# Fire effect and its legacy modulate soil bacterial and fungal communities in Chinese boreal forests along a chronosequence

**DOI:** 10.1101/2020.07.31.231910

**Authors:** Wei-qin Su, Caixian Tang, Jiahui Lin, Mengjie Yu, Yu Luo, Yong Li, Zhongmin Dai, Jianming Xu

## Abstract

Wildfire has increasingly profound and pervasive consequences for forest ecosystems via directly altering soil physicochemical properties and modulating microbial community. In this study, we examined the changes in soil properties and microbial community at different periods after highly severe wildfire events (44 plots, 113 samples) in the Chinese Great Khingan Mountains. We also separated charcoals from burnt soils to establish the relationship between soil microbes and the microbes colonized on the charcoal. Wildfire significantly altered bacterial and fungal community structures across a 29-year chronosequence. The network analysis revealed that from 17 years after fire, the complexity and connectivity of bacterial and fungal communities were significantly increased. Differential abundance analysis suggested that bacterial and fungal OTUs were enriched or depleted only during 0-4 years after fire. In addition, soil factors, including soil pH, total C and N, soil water content, and dissolved C and N, are key determinants of soil bacterial and fungal communities from 17 years after fire. The fire-derived charcoals provided a new and unusual niche for microbial colonization and charcoal microbes had a significantly different community structure from the burnt soil microbes. Our data suggest that soil bacterial and fungal communities changed dramatically during the recovery from fire events in terms of the abundance and co-occurrence networks in the boreal forest ecosystems.

**Importance:** Pervious research has reported fire altered soil microbial community composition and function during short-term succession in boreal forests. However, the long-term effect of fire and fire-derived charcoals which are regarded as fire legacy effect on soil bacterial and fungal communities composition and structure have not previously been shown. Understanding how soil microbes particularly the keystone taxa and determinative soil factors, respond to fire and its legacy matter charcoal, is critical for predicting how future fire influences soil nutrient transformations and biological processes. We accessed time chronosequence to examine the effect of fire history on soil microbial abundance and co-occurrence network. These findings suggest that soil microbes can be reshaped by fire and its legacy effect of fire-derived charcoal even in the long periods after fire and provide further insights into fire and its legacy effect.

## Introduction

Fire is one of the most important disturbance agents in terrestrial ecosystems and a worldwide phenomenon in our earth (1). Ecosystems such as boreal forests, shrublands, grasslands and savannas, which have been regarded as “flammable systems”, are often disturbed by wildfires over millions of years (2). Triggered by heat, fire affects soil microbiota and nutrient cycling directly and indirectly (3, 4). Basically, severe wildfire changes critical biotic and abiotic processes to cause complex consequences, including significant removal of plant shoot and organic matter, destruction of the soil physical structure and porosity, increase in nutrient losses through leaching and volatilization and considerable shift of chemical properties (3, 5–7). Fire breaks biomolecules (8) by exposing soil microbes directly to extremely high temperatures and lowers bacterial and fungal biomass (by 33 - 49%) (4, 9). Meanwhile, fire-based disturbance also modifies microbial community diversity and phylogenetic structure, which link to carbon dynamics and thawing of permafrost (10, 11).

Boreal forest is a major fire-prone biome, covering approximately 30% area of the global forest area. Distributed in high-latitude regions of Eurasia and North America, boreal forests have low productivity and are easily forced by fire and climate change (12). During long-term persistence and repeated succession, boreal forest ecosystems are tied to corresponding fire patterns and post-fire effects. Future climate change (such as global warming and drought) is likely to increase fire frequency and severity (13–15). Fire regimes have brought international concerns in the global boreal forests.

In China, boreal forest landscapes encompass the most southern parts of the global boreal forest (16). Most fires occur as a surface fire which is usually of moderate to high severity, with an estimated historical fire interval period of 30-120 years (17). Chinese boreal forests store approximately 24-31% of total carbon storage (1.0-1.5 Pg C) in China (14) and the average of annual carbon emissions from boreal forest fire up to 0.54 Tg (18). Moreover, strict fire suppression has been implemented in this region for over a half of century (19), since a catastrophic wildfire (20) occurred on 6 May 1987. Such a human control has already changed the fire regime—the patterns of fire spread, intensity, severity, frequency, seasonality and ecological effects (2, 21).

The formation of charcoals is another important phenomenon during vegetation burning, especially in the boreal forest region. Most carbon is converted to CO_2_ releasing to the atmosphere while a small portion of forest fuel (plant biomass and soil organic matter) is pyrolyzed into other forms of pyrogenic carbon (PyC) (22). The PyC is a term synonymous with black carbon (23) and is mainly produced as solid charred residue including visually-defined charcoal and much lower proportions of volatile soot (24). Charcoal is part of the PyC continuum. Because of its resistance to decomposition, it serves as (1) historic records of fire and estimates for fire activities (1, 25), (2) an important long-term carbon sink (22, 23, 26) and (3) different habitat patches for soil microorganisms (27–29). In addition, incorporation of fire-derived charcoal into forest soils has an impact on soil C and nutrient cycles, and vegetation regeneration (30–32). Owing to the presence of charcoal, burning *per se* contributes to the shifts in ecological properties (including soil biological properties). Therefore, charcoal is thought to function as fire’s black legacy. However, some studies have revealed that charcoal produced in fire would re-mineralize or degrade more rapidly than expected (33, 34). Charcoal persistence directly influences soil properties in decades and indirectly affect microbial community succession.

The effect of fire on soil biogeochemical and microbial processes can last for decades and even centuries (35). The short-term effects of fire on soil pH, water content, organic matter content, soil autotrophic respiration, and concentrations of total N, ammonium and cations (K, Na, Ca, and Mg) are also observed in the boreal region (7, 37, 38). These effects on soil properties in turn influence soil microbial community diversity and structure (7). However, recovery patterns of soil microbial community and the functioning from fire events are currently unclear in boreal ecosystems. The information is limited about the direct and indirect impacts of fire, associated changes in edaphic factors and fire-derived charcoal on soil microbes over long-term scales. The knowledge gap still exists in understanding complex and diversified interactions among soil microorganisms with the soil environment (such as in the soil-charcoal system). In addition, we use “space-for-time” substitution (39) as a judicious strategy for chronosequence research. On the other hand, it is difficult to separate charcoal particles from the soil and hence to identify specific taxa colonized in the charcoal or soil samples. Consequently, we are not able to quantitatively assess the effect of each individual factor even if charcoal could affect microbial community assembly through several known pathways (40).

In this field study, we used a 29-year chronosequence to examine the effect of fire history on soil microbial community recovery processes. Specifically, we quantified the responses of soil microbes to fire and its legacy effect of fire-derived charcoal, and determined the key determinants of bacterial and fungal community structure. We hypothesized that (i) bacterial and fungal co-occurrence networks would become more complex and connected with increasing time since fire (TSF), and (ii) the fire-derived charcoal provided a unique niche for soil microbes and had microbial communities distinct from the soil, and (iii) some soil properties would be the key determinants of soil microbial community structure in soils with different fire histories.

## Results

### Effect of fire on soil properties and microbial communities

Fire enhanced the mean concentration of soil available P from 5.4 to 19.3 μg g^-1^ (*p* < 0.01) (Table S2) in A horizon. Compared with the unburnt controls, middle-term fires (MF) increased available P by 224% and old fires (OF) increased available P by 257%. However, no significant changes (*p* > 0.05) in soil properties were observed in O horizon. Severe wildfires significantly increased soil C/N ratio and the concentrations of available P in both horizons (*p* < 0.05) (Fig. S2). There was no statistical difference in other soil properties (Fig. S2).

In total, 10480 bacterial and 56 archaeal OTUs were identified from 16S sequences, while 2711 classified fungal OTUs assigned from 18S sequences. Proteobacteria (30.6%-40.4%) were the most abundant phylum of bacteria overall in the soil samples (Fig. 1d), followed by Actinobacteria (22.2%-32.0%), Acidobacteria (12.6%-16.6%) and Verrucomicrobia (4.2%-10.9%) (Table S3). Relative abundances of Proteobacteria, Actinobacteria and Acidobacteria were generally constant across different fire histories (Table S3), irrespective of soil horizon. Most of fungal OTUs were in Ascomycota (41.3%-60.1%), with fewer in Basidiomycota (26.1%-49.4%) and Mucoromycota (8.3%-13.2%) (Table S3). The proportions of dominant phyla were variable in the post-fire soils. Overall, there is no significant temporal variation on taxonomic composition after fire (Fig. 1d). Alpha diversity indices (includes Shannon index and OTU richness) of bacterial and fungal communities did not vary significantly between the fire-affected (RF, MF, OF) and unaffected groups (UF) (Fig. 1c). However, wildfire caused significant differences (*p* < 0.001, PERMANOVA) in β-diversity of soil microbial community (Fig. 1a & b) across different fire histories (RF, MF, OF and UF). The NMDS of Bray-Curtis dissimilarities also displayed that bacterial and fungal β-diversities in O horizon were significantly (*p* < 0.005, PERMANOVA) different from those in A horizon (Table S4).

**Figure 1.**
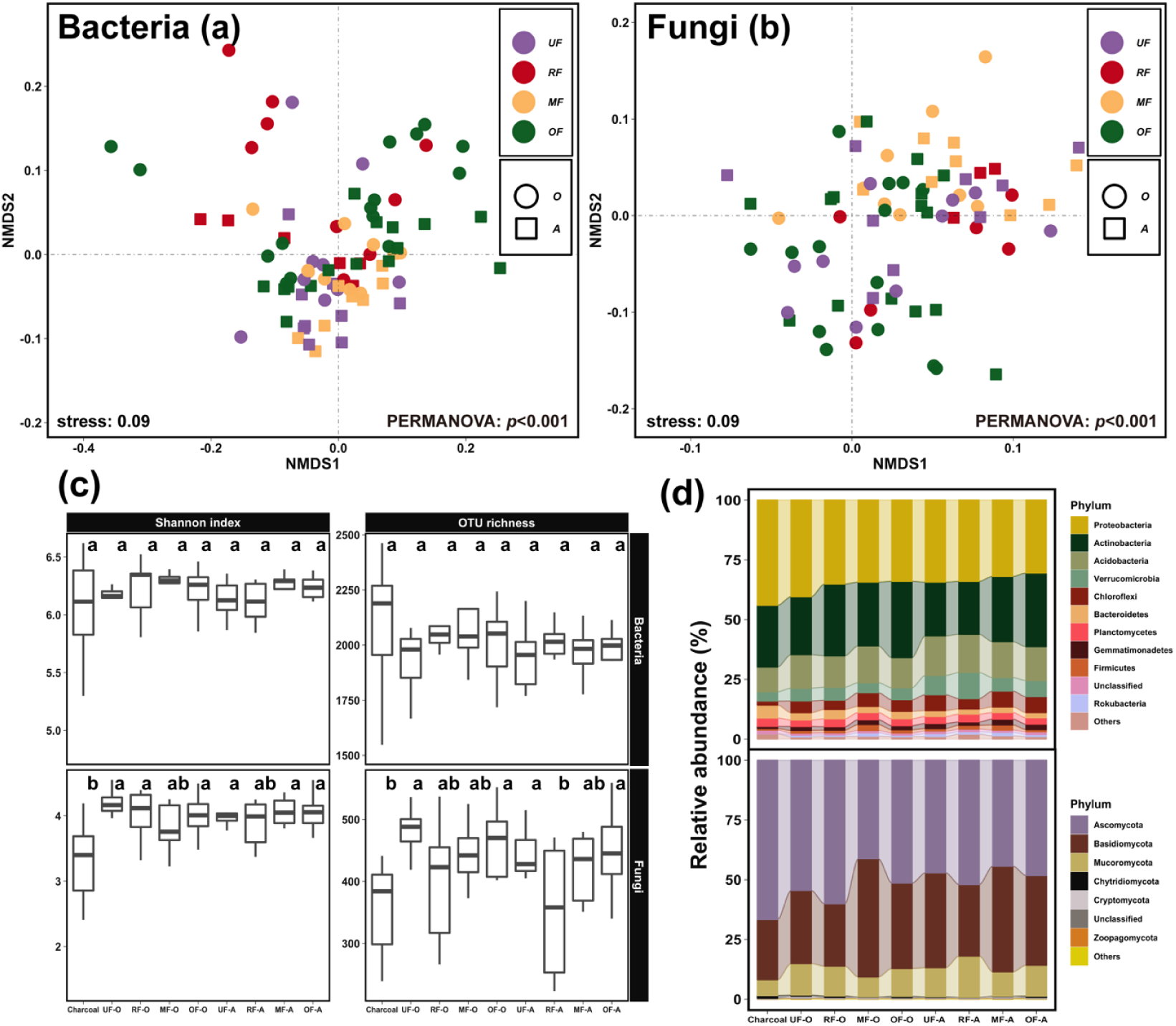
Microbial composition and structure in soils and charcoal samples of different fire histories and soil horizons. Non-metric multidimensional scaling ordination plot based on Bray-Curtis dissimilarity showing the change of soil bacterial (**a**) and fungal (**b**) community structure. Purple, red, orange and green symbols represent unburnt controls (UF), and recent (RF, 0-4 years), middle-term (MF, 7-15 years) and old fire (OF, 17-29 years), respectively. As listed in the legend, solid circles and squares represent samples from the O and A horizons, respectively. Boxplots of *α*-diversity (**c**) for bacterial and fungal communities in the soils and charcoal samples of different fire histories (UF, RF, MF and OF) in O and A horizons. Boxes are bounded on the first and third quartiles, divided by median lines. Boxes with different lower-case letters are significantly different (*p* < 0.05) by Bonferroni’s post hoc tests. Relative abundances of bacteria and fungi among different fire histories in O and A horizons at the phylum level (**d**). “Unclassified” OTUs indicate no taxonomic information matched in the database. Charcoal samples only were separated from burnt soils in O horizons.

### Effect of fire on differential OTUs

Differential abundances in bacterial and fungal community composition after wildfire across a chronosequence were identified (Fig. 2). The enrichment and depletion effects on bacterial communities in O (20 vs. 0) and A (13 vs. 1) horizons were the most remarkable in the recent burnt soils (RF). There are no differential abundances in bacterial community composition after middle-term and old fires, compared with the unburnt soil (Table S5). After fire (RF vs. UF), the changes of bacterial composition were mostly positive. In the period of 0-4 years after fire, differential OTUs had at least 3.5-fold enrichment in bacteria in both O and A horizons (Fig. 2a). There were a number of spore-forming bacterial taxa, including the group of Terrabacteria (Battistuzzi and Hedges, 2009). Terrabacterial phyla (Order Bacillales and Order Micromonosporales) were only significant in O horizon but not found in A horizon. However, the abundance of many fungal OTUs (e.g. Ascomycota and Basidiomycota) were declined significantly after recent fires, especially in A horizon (Fig. 2b). Similarly, the RF group also had the most enriched and depleted fungal OTUs in O (17 vs. 7) and A (11 vs. 20) horizons compared with the unburnt soils (Table S6). There were more fungal differential OTUs in A horizon than in O horizon (Fig. 2b). Moreover, the MF group had only one significantly enriched OTU while the OF group had only one depleted OTU (Table S6).

**Figure 2.**
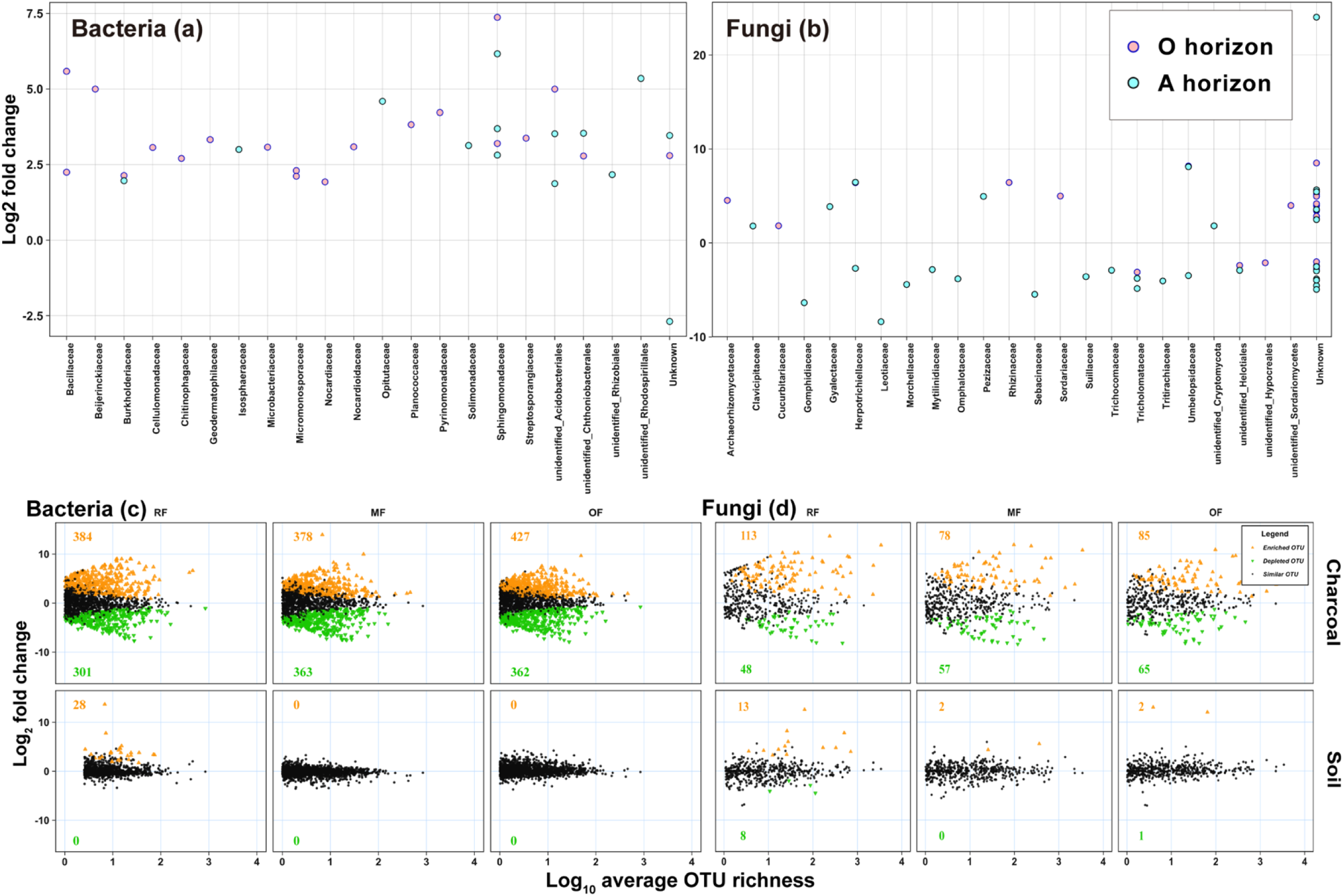
Enrichment (above 0) and depletion (below 0) of bacterial (**a**) and fungal (**b**) OTUs in soils after recent fires (RF, 0-4 years) compared with unburnt soils by differential abundance analysis. The OTUs (singleton is exclusive) are arranged by family on the x-axis and each point represents an individual OTU. The y-axis indicates fold change in log base 2 units. Skyblue points represent differential OTUs found in O horizon while pink points represent differential OTUs found in A horizon. The significant threshold is 0.01. Enrichment (yellow triangles) and depletion (green triangles) of OTUs (singleton is exclusive) about bacterial community (**c**) and fungal community (**d**) by differential abundance analysis. Enrichment and depletion of certain OTUs in the charcoal and soil samples of recent (RF, 0-4 years), middle-term (MF, 7-15 years) and old fire (OF, 17-29 years) compared to unburnt soil samples (black dots) in O horizon. The significant threshold is 0.01. Colored numbers represent the number of differential OTUs.

### Bacterial and fungal co-occurrence networks

To identify the co-occurrence patterns of bacterial and fungal communities in soils with different fire histories, we constructed bacterial and fungal networks for the two horizons. The networks displayed remarkable differences in their structure and topology (Figs 3 & 4). The bacterial network of the UF group in O horizon consisted of 64 nodes (i.e. taxa) and 57 edges (associations between taxa), while the RF group in O horizon consisted of 60 nodes and 30 edges. In stark contrast, the OF group in O horizon consisted of 222 nodes and 380 edges (Table 1). The average degree and the average number of neighbors of bacterial network in the OF group were also considerably higher than the UF and other fire-affected groups. The higher complexity and connectivity in the OF bacterial network showed that the special and complicate modules had been formed during long periods after wildfire (Fig. 2d). There were similar results in A horizon that more complex coupling among bacteria occurred during the post-fire succession process (Figs S5a & d).

**Table 1.**
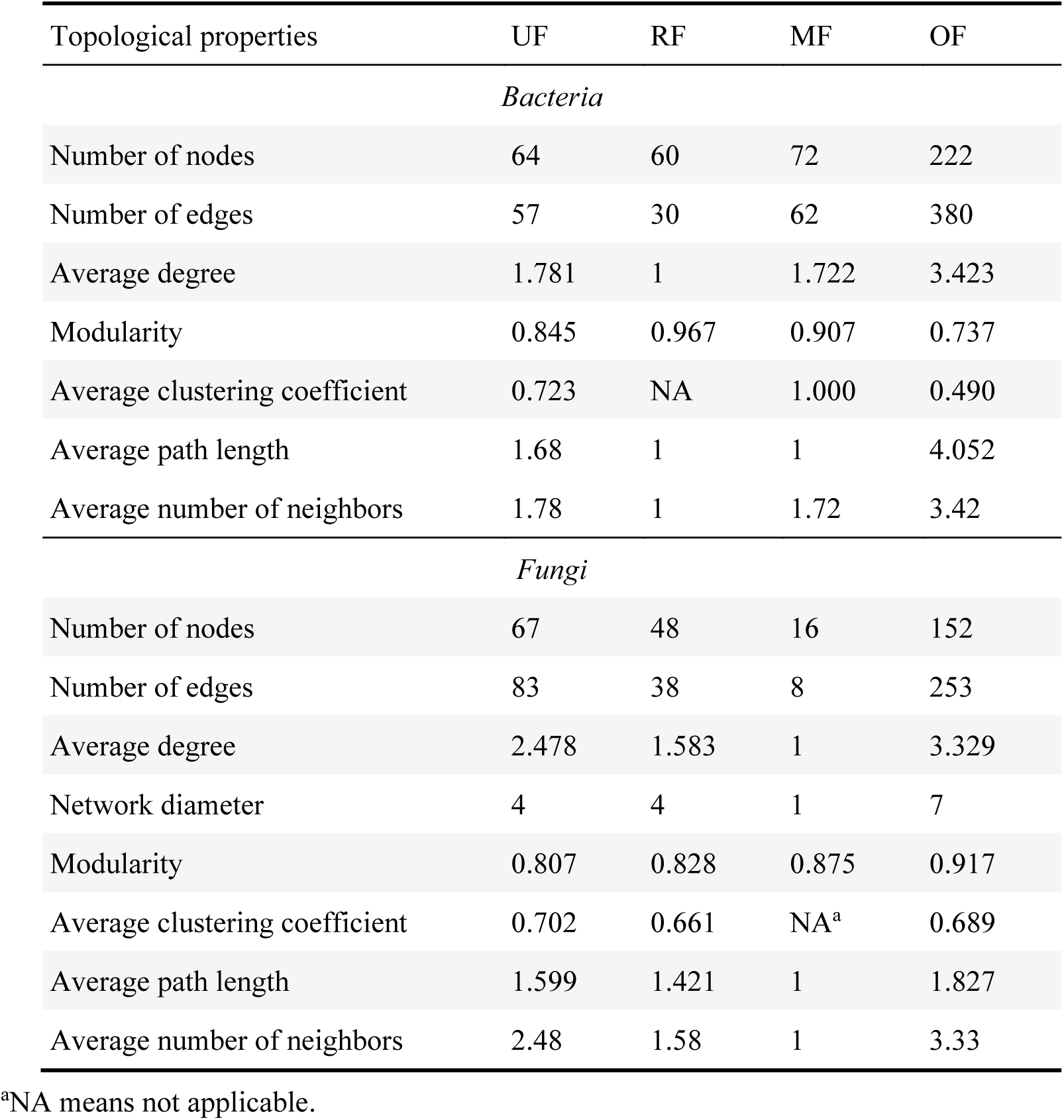
Topological properties of networks obtained at the different fire histories in O horizon. Fire histories are recent fires (RF), middle-term fires (MF), old fires (OF) and unfired (UF).

**Figure 3.**
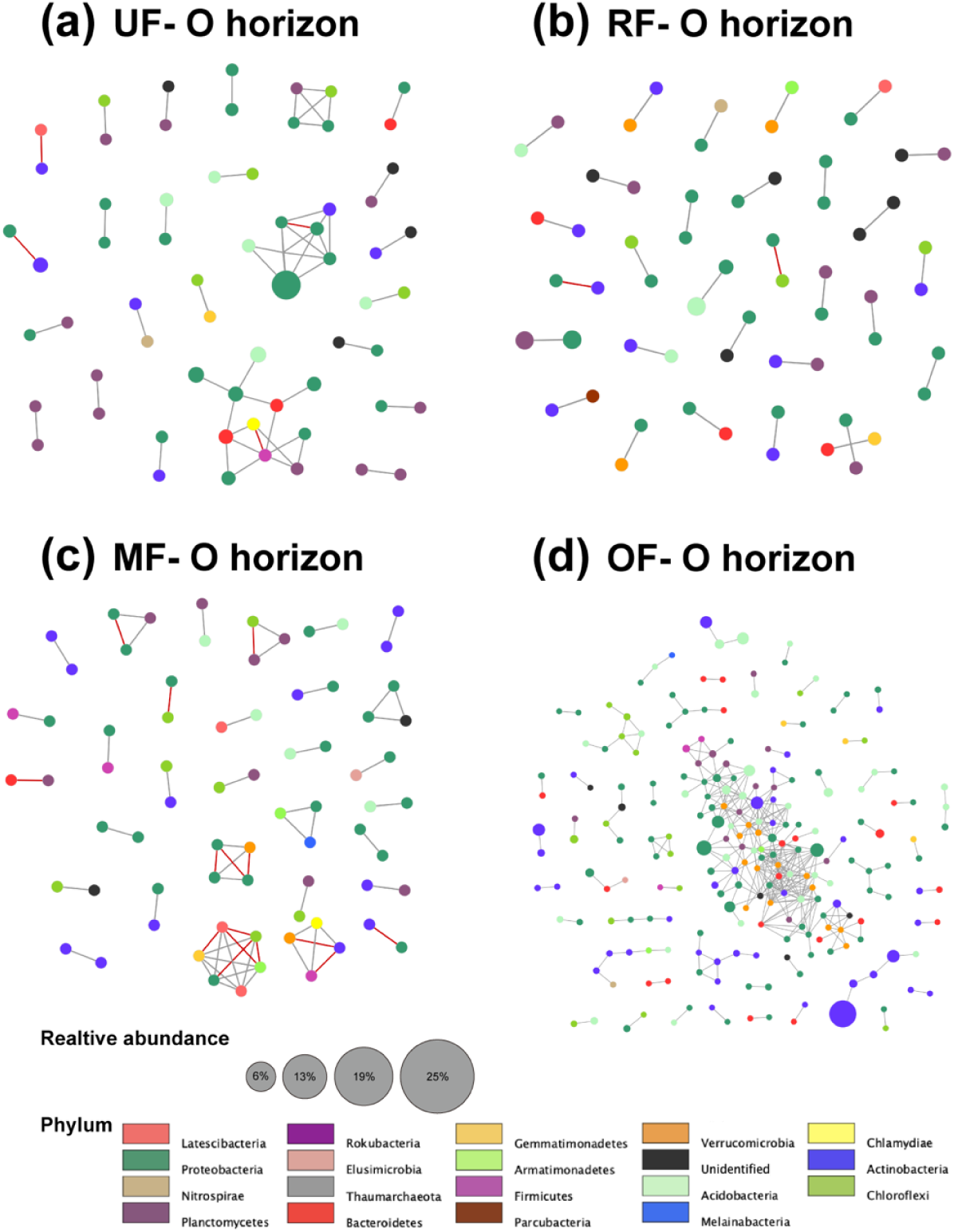
The co-occurrence networks of the bacterial community in O horizon under recent fires (RF, **b**), middle-term fires (MF, **c**), old fires (OF, **d**) and unburnt groups (UF, **a**). A connection stands for a strong (Spearman’s *p* > 0.8) and significant (*p* < 0.01) correlation. The size of each node is proportional to the relative abundance, as is shown in the legend. The thickness of each edge is proportional to the value of Spearman’s correlation coefficients. The grey edges represent positive interactions between two bacterial nodes, while red edges represent negative interactions.

**Figure 4.**
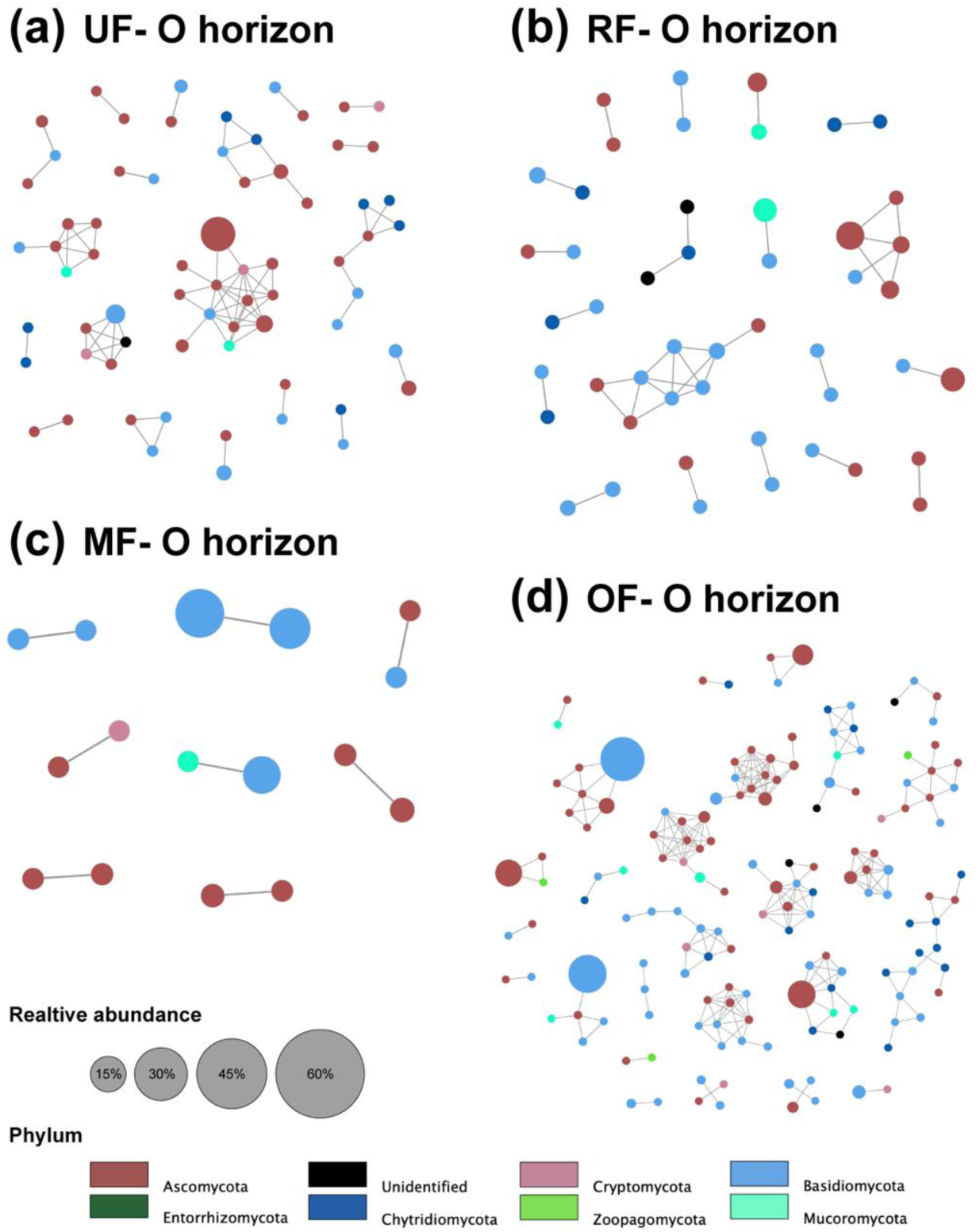
The co-occurrence networks of the fungal community in O horizon under recent fires (RF, **b**), middle-term fires (MF, **c**), old fires (OF, **d**) and unburnt groups (UF, **a**). A connection stands for a strong (Spearman’s *p* > 0.8) and significant (*p* < 0.01) correlation. The size of each node is proportional to the relative abundance, as is shown in the legend. And the thickness of each edge is proportional to the value of Spearman’s correlation coefficients. All edges are grey, which means positive interactions between two fungal nodes.

Multiple topological properties of the fungal co-occurrence networks pronouncedly varied in different fire histories (Table S7). The fungal network of the OF was largest with 152 nodes and 253 edges and had the highest average degree in O horizon (Table 1). Higher complexity in the OF group was visible, indicating that many fungi had developed a number of associations (Fig. 3). We observed that both modularity and average path length of fungal network in the OF group were highest in O horizon, showing more structured fungal communities within the network. Unlike bacterial patterns, fungal co-occurrence patterns of the OF group were more modular than the corresponding unburnt soil networks. Moreover, the connections between fungi in A horizon were gradually increased after wildfire disturbance (Fig. S6).

### The effects of fire-derived charcoal

To gain insight into the fire legacy effect, we compared microbial properties in charcoal samples with soils. Despite common patterns in the relative abundance of main taxa at the phylum level, microbial community composition, especially for fungi, had some apparent divergences between charcoal and soil, and between different fire histories (Fig. 1d). The predominant phyla (ascomycota, basidiomycota, and mucoromycota) significantly differed in their relative abundances in the charcoal and unburnt soils (Table S3). Unlike burnt soils, charcoal samples had an enhanced colonization of differential bacterial and fungal OTUs from the unburnt controls (Fig. 2c & d). On the other hand, the burnt soil samples of O horizon had more similar microbial composition to the unburnt sites than to charcoal samples, as indicated by nearly zero differential OTUs (Fig. 2c & d). Bray-Curtis dissimilarity was used to assess the structure of microbial community colonized on the surface of fire-derived charcoal or inhabited within the soil. NMDS ordination revealed significant differences (*p* < 0.001, PERMANOVA) among charcoal, unburnt, and burnt soil samples (Fig. S4) for bacteria (R^2^ = 0.30, *pseudo* F = 23.5) and fungi (R^2^ = 0.19, *pseudo* F = 13.2). These results showed that the charcoal colonized distinct microbial community, as fire legacy effect. The explanation of variance to 40.7% in the soil fungal community structure on SEM (including charcoal microbial community) while explanation of variance decreased slightly to 41.4% for bacterial community structure (Fig. 6a & b). Based on SEM, fire significantly affected bacterial and fungal community in charcoal and bacterial community in charcoal also significantly affected bacterial community in soil.

**Figure 5.**
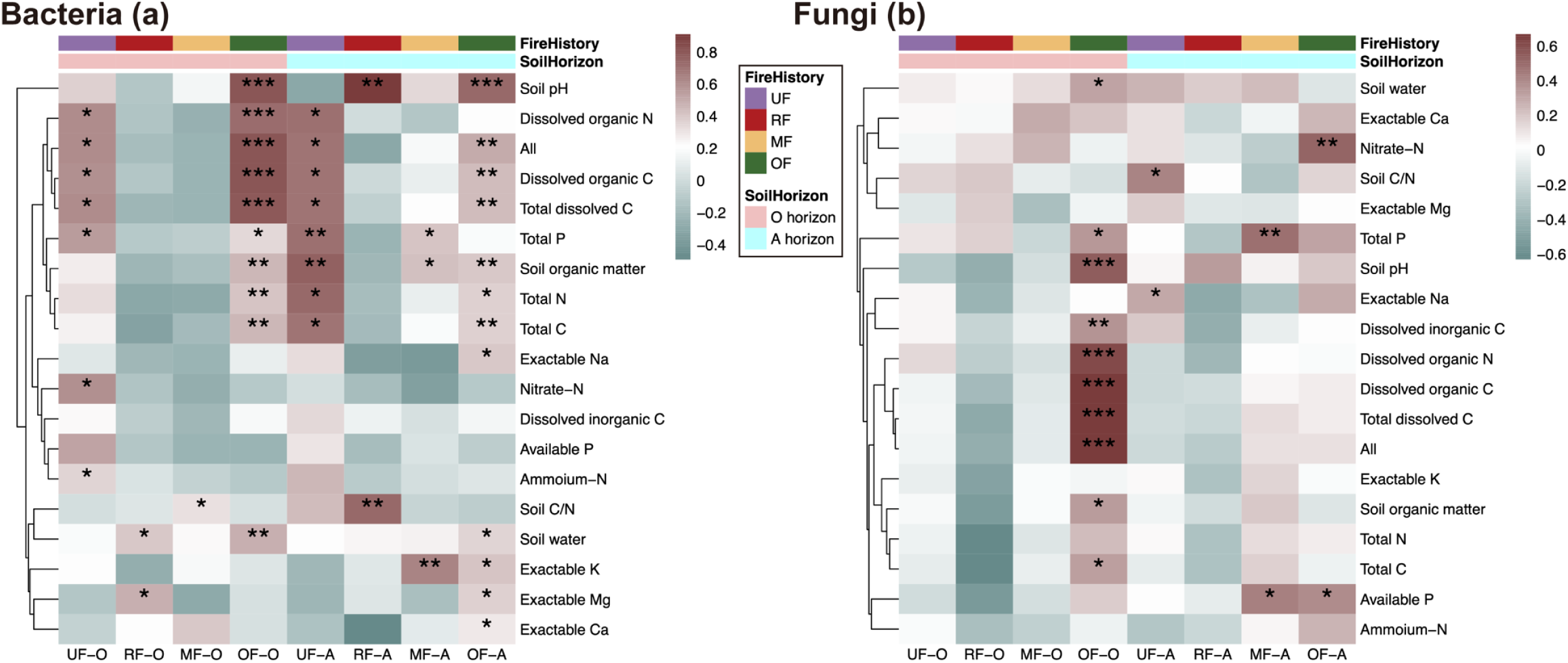
Associations of bacterial (**a**) and fungal (**b**) community structures with soil factors, by Partial Mental test, in different fire histories and different horizons (O and A). Fire histories are recent fires (RF), middle-term fires (MF), old fires (OF) and unfired (UF). All, all environmental factors. The correlations (r) and significance (*p*) are determined with a Bray-Curtis distance matrix and a Euclidean distance matrix for environmental factors, controlled with spatial variation. Different colors in the cells indicate different mantel statistic correlations (r), shown in the legend. The *p* values in the cells with *, ** and *** indicate *p* < 0.05, *p* < 0.01, *p* < 0.001, respectively. Significance for each test is calculated from 9999 randomized Monte Carlo runs.

**Figure 6.**
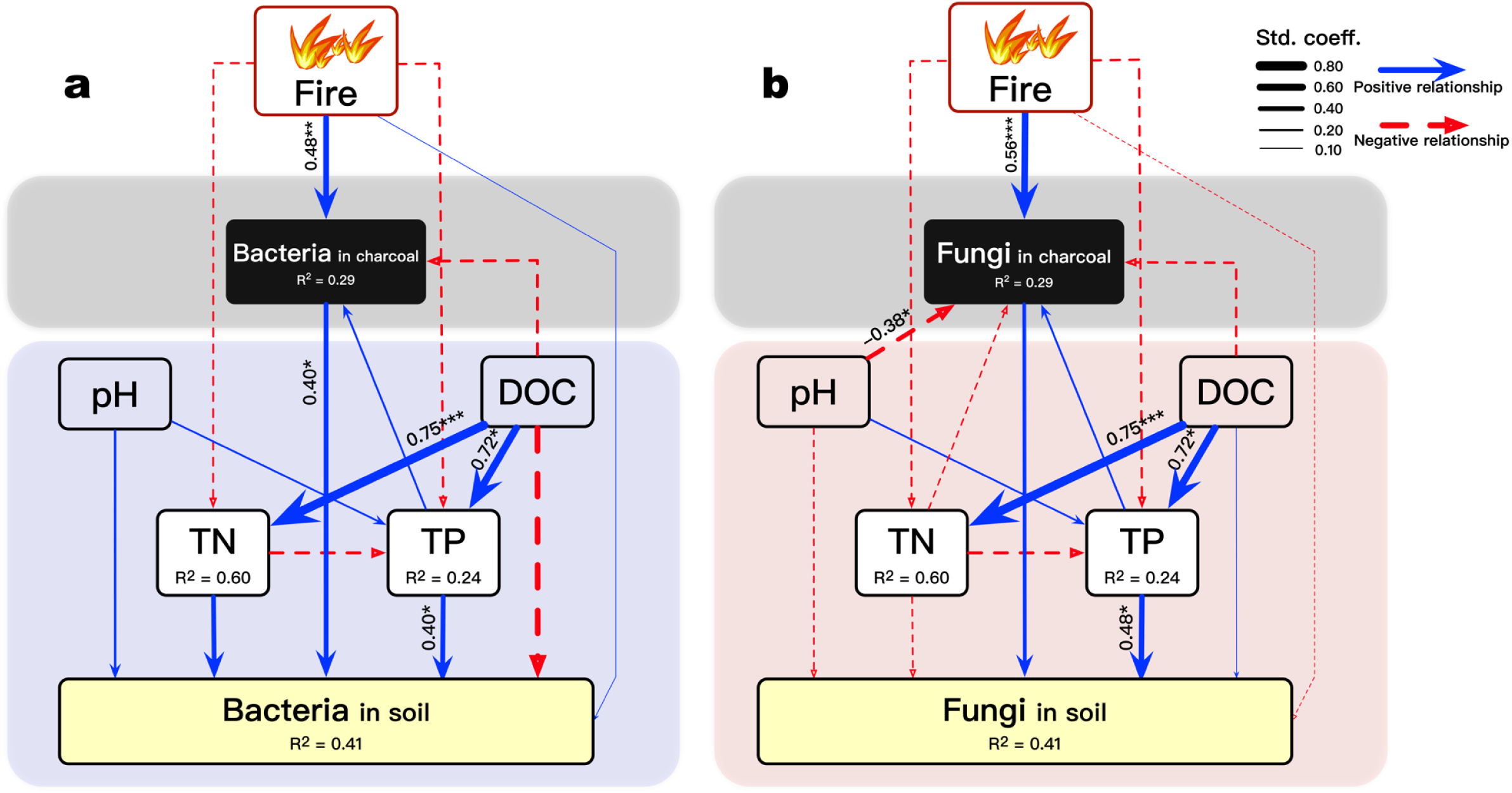
Fire legacy effect on bacterial (**a**) and fungal (**b**) community structures of the burnt soil in O horizon. Best-fitted structural equation models depicting the major pathways of fire, charcoal and representative environmental variables on microbial community structures. The bacterial and fungal community structures of charcoal and soil represented by the first axis of NMDS. DOC, dissolved organic carbon; TN, total nitrogen content; TP, total phosphorus content; Fire, time since fire. R^2^ values denote the amount of variance explained by the model. Solid (blue) and dashed arrows (red) indicate positive and negative effects, respectively. Arrow thickness is proportional to the standardized coefficients (std. coeff.), interpreted as the relative importance of effects. Only significant (\**p* < 0.05, \*\**p* < 0.01, \*\*\**p* < 0.001) estimate numbers are noted at each arrow. The goodness of fit was satisfactory (n = 31, *p* value = 0.483, CFI = 1.000, RMSEA = 0.000, SRMR < 0.015).

### The influence of soil properties on microbial community

The relationships between soil properties and soil microbial community structure varied with different fire histories and horizons (Fig. 5). Soil pH, soil organic matter, total N, total P, dissolved organic C, and dissolved organic N had significant relationships with bacterial community structure in both O and A horizons of the OF group. However, in the RF and MF groups, most of soil factors had no significant correlations with bacterial community structure. Soil factors, such as pH, soil organic matter, total P, dissolved organic C and dissolved organic N, significantly correlated with the structures of fungal community in O horizon (Fig. 5). More soil factors had significant relationships with bacterial than fungal community structure. In O horizon of the OF group, bacterial community structure correlated highest with pH (r = 0.8095, *p* = 0.0001) while fungal community structure correlated highest with dissolved organic C (r = 0.6691, *p* = 0.0001) across all the environmental variables (Table S8).

## Discussion

### Temporal patterns of bacterial and fungal communities in response to fire

This study demonstrates that wildfire has an impact on bacterial and fungal composition, and thereby modulates the patterns of soil microbial community succession over three decades after fire in the boreal forest region. Translated spatial differences into temporal changes, our results clearly show that temporal patterns of bacterial and fungal communities occur across a chronosequence. Supported by previous findings about bacterial diversity shifts under fire disturbance (7), wildfire does not show significant impacts on microbial *α*-diversity (Shannon index and OTU richness) both in O and A horizons. Only fungal *α*-diversity in A horizon significantly declined by the recent fires, corresponding with fungal co-occurrence networks in this horizon (Fig. S6b). This is in agreement with previous studies that fungal diversity changed significantly in 2 years after fire across a 152-year chronosequence (41). In addition, the fire-related effect on species richness of ectomycorrhizal fungi has not been observed on many fire-affected soils, and mycorrhizal colonization of roots may increase after a fire event (42). However, in Alaskan boreal forests, soil fungal community diversity (Shannon index) varied significantly only in the 90-year-old site, not in sites with shorter-term fire history across a 100-year chronosequence (43). Therefore, soil bacterial and fungal community diversities have concomitantly and rapidly recovered, suggesting that fire only has a limited impact on bacterial and fungal diversity across a chronosequence.

In general, wildfire is reported to have a distinct effect on soil microbial community composition. Our results demonstrate that the bacterial and fungal community compositions differ among fire history. Bacterial and fungal community structures have an underlying trend clustered with increasing TSF (Fig. S3), which means fire has a significant effect on soil microbial structure across a chronosequence. This phenomenon goes beyond previous results reporting that bacterial community altered dramatically after 16 weeks since fire (44) and could return to similar levels of the controls after 11 years (45). Yet, there are different temporal patterns for bacteria and fungi under fire disturbance. Bacteria under the MF and UF groups were closely clustered in the NMDS plot. Their co-occurrence networks and topological networks are similar to those of other fire-affected groups in both O and A horizons (Fig. S5). However, fungal co-occurrence always differs between the fire-affected groups and the unburnt group, suggesting that the fungal communities still change after 29 years since fire. Our partial Mantel data show that bacterial community structure is correlated with many soil factors in the OF group but correlated with few soil factors in the RF and MF groups. These results indicate that fire-derived changes in soil properties become the major determinants of soil microbial community composition in the post-fire succession process. Our analysis further suggests that soil properties have significant effects on soil microbial community especially for bacterial community as environmental filters from 17 years after fire. Furthermore, bacterial community was complexly modulated by soil factors more significantly than fungal community.

Soil microbial co-occurrence networks, that provide insight into ecological interactions among taxa, were influenced by fire history. The major changes in topological properties of the bacterial and fungal networks have more nodes and edges under the OF group than the UF group (Figs 2, 3, S5 & S6), predicting that microbial communities have more connections and complex functional modules under fire disturbance. Fire reshapes soil microbial network and makes microbial interaction stronger in both O and A horizons after 17 years since fire. In addition, during 1-15 years since fire (RF and MF groups), taxa only form weak relationships and lower-complexity modules in most of the samples. Clearly, bacteria and fungi need more than 15 years to recover and reconstruct ecological functioning after fire events. Our results are in contrast to the short-term experiments (within 12 months after fire) showing that fire did not alter co-occurrence network generally (46).

### Specific bacterial and fungal taxa in response to fire

By using differential OTU abundance analysis across a chronosequence, we observed that only RF groups serve an enrichment/depletion role for microbial OTUs relative to the unburnt soil. There are few differential OTUs found in the MF and OF groups although microbial co-occurrence network in the MF and OF groups is significantly different from that in the UF group. These results indicate the number of most OTUs in post-fire boreal soil back to pre-fire level after 7 years. Many bacterial species are able to cope with extreme environments such as high temperature by forming resistant substances such as spores and endospores (10). We observed that the RF group serves an enrichment role for a subset of bacterial OTUs relative to the unburnt soil. The recent fires increased the number of spore-forming OTUs in the order Micromonosporales (phylum Actinobacteria) and the Gram-positive Bacillales (phylum Firmicutes). The Bacillales can generate dormant and resistant spores that make them survive under harsh conditions for years (47). The Bacillales, that consist of many spore-formers, allow themselves to recover better and faster from stresses. Micromonosporales (genus Actinoplanes) also form spores by fragmentation to reproduce (48). These post-fire increases in the number of bacterial OTUs are more in O horizon than in A horizon, which could be linked to higher fire severity or upper fire loading density in O horizon. The results of these differential bacterial OTUs are consistent with the findings of many previous studies showing pronounced effects on soil bacterial community in the short-term, mostly within 36 months after fire (6, 10, 49). Furthermore, apart from fire severity, environmental factors (soil pH, soil water content, total C, dissolved C and N) are also key determinants driving soil bacterial community. Some bacterial OTUs depend on plant-soil feedback (50) and are strongly affected by aboveground community during forest regeneration (51, 52).

We observed that there are many depleted ectomycorrhizal OTUs (such as Family Tricholomataceae) in O and A horizons, which is consistent with a previous study showing that forest fire caused a significant loss of ectomycorrhizal fungi biomass in the litter and organic horizons (53). Severe fire consumes the most portion of humus on soil surface and kills most of trees, and hence the death of ectomycorrhizal fungi although some ectomycorrhizal fungi have sporophytes (54). Moreover, during the early successional phase after fire disturbance, the regeneration has not reached pre-wildfire levels in the Great Khingan Mountain. The impact of vegetative recovery on fungal community may play an important role in enriched OTUs in O horizon. Furthermore, most of differential OTUs in A horizon are depleted probably because vegetation including herbaceous species and shrubs still cannot influence soil microbes in A horizon. However, many fungal species can still survive and largely remain intact even after a high-intensity wildfire. The regeneration of post-fire pyrophilous ascomycetes is well documented for the reasons such as the tolerance of high pH and other physicochemical consequences of wildfire (42).

### Fire legacy effect on soil microbial community

Our study, for the first time, examined the effect of fire-derived natural charcoal on soil microbial community along a chronosequence in a boreal forest region, and showed that the microbial community colonized on the charcoal can influence the responses of soil microbial community to wildfire. Due to specific porous structure and absorbed nutrients, charcoal is a habitat for soil microbes and these microbes are significantly different from those in the burnt soils (Fig. S4). The major taxa and *α*-diversity of microbes in the charcoal (Figs 1 & S4) are also distinct with the burnt forest soils, highlighting a new and unique habitat produced by fire (28). Moreover, there are more than 680 bacterial OTUs and more than 120 fungal OTUs in the charcoal samples (Fig. 2c & d). Together these observations provide strong evidence that fire-derived charcoal create distinct environments (including abiotic factors and biotic factors) as the habitat for fire-responsive microbes. Because of its higher pH value, charcoal causes a localized increase in soil pH (55, 56) and shifts the way of soil pH effect on soil microbial community to some extent. Charcoal microbial community has a significantly positive relationship with soil bacterial community (Fig. 6a), indicating that microbes in soil are more likely impacted by post-fire charcoal. Charcoal microbial community is also related to soil pH, dissolved organic C and total P which regulate microbial activity with nutrient cycling. Previous studies have shown that charcoal absorbs some labile C compounds to inhibit or activate some microbial processes (29, 40, 57). However, our present study has not found that dissolved organic C has any specific relationship with bacteria or fungi in the charcoal. Based on SEM, fire (represented by TSF) affects soil microbial community via the indirect effect of charcoal. Furthermore, charcoal community structure varied with TSF showing that the ecological recovery processes after fire occur in the charcoal particles and the fire legacy effect gradually changes over time.

### Conclusions

This study provides the key evidence how large-scale wildfire events and the derived charcoal (fire legacy effect) affect soil microbial community along a 29-year chronosequence of the boreal forest region in northeast China. It reveals that wildfire significantly impacts on bacterial and fungal community structures. The complexity and connectivity of bacterial and fungal communities were significantly enhanced from 17 years after fire when soil pH, total C and N, soil water content, and dissolved C and N are key determinants of soil bacterial and fungal communities. Furthermore, the charcoal formed at fire events and its colonized microbes have important ecological functions of mediating post-fire successional boreal forests although our current knowledge of the succession patterns about soil microbes post-fire and charcoal functions in flammable boreal forest ecosystems is limited. Our research highlights an important step forward to clarifying the effects of fire and its charcoal on soil microbial community. Further research may aim at (i) identifying specific physiological and biochemical processes in response to fire events and fire history and (ii) distinguishing charcoal microbial community from post-fire soil microbial community in finer details.

## Materials and methods

### Study area and experimental design

This study was carried out in the boreal forest region, the Great Khingan Mountains (50’10’N-53’33’N, 121’12’E-127’00’E), north-eastern China (Fig. S1). The mean annual air temperature in this region is -2 to -4 °C and the mean annual rainfall is 350∼500 mm (18). The soil at the sampling sites is classified as dark brown forest soil (Inceptisol) and the parent material is granite bedrock (45, 58). The vegetation of this region is representative of boreal coniferous forests, forming the southern extension of the Siberian boreal forests. Transition plant species, dominated by larch (*Larix gmelini*), are late-successional and widely distributed from a wildfire (14) or harvesting. Broadleaf trees as the pioneer species, mainly birch (*Betula platyphylla*), are the earliest re-generation in the post-fire soil. They are succeeded by boreal conifer tree species in the wildfire-disturbed boreal forest ecosystems.

We selected a total number of 44 plots (50 × 50 m) at altitudes ranging from 209 m to 466 m a.s.l. (Table S1) in July 2015. Plots including fire-affected and unaffected groups, were located at the National Reserve and State-owned Forest with minimal human interventions (e.g. prescribed fire, and logging). Due to the difficulty of long-term studies and specificity of fire researches, we followed a classical space-for-time substitution method to assess the impacts of fire and its legacy charcoal. Fire-affected groups consisted of a chronosequence of burnt forests that were representative of different TSF (59), considered as different fire histories. These affected groups were divided into Recent Fires (RF: 0-4 years since fire, n=9), Middle-term Fires (MF: 7-15 years since fire, n=12), and Old Fires (OF: 17-29 years since fire, n=12) based on TSF. Each fire-affected site was considered as an independent replicate and more information about TSF is given in the Supplementary Information.

According to the historical records from the State Forest Administration and remote sensing images, each of the fire-affected sampling sites had suffered one highly severe wildfire during 1915-2015. The highly severe wildfire here means destroying at least one hundred hectares of forests and removing litter and organic soil layer (60). To save expenditure, we only sampled fire-unaffected controls (UF, n=11) nearby the burnt plots with distinct burnt borders. The minimum distance between each plot was 100 meters. For reducing spatial heterogeneity, all plot samples were comprised of more than five subsamples (10 × 10 m) and each subsample was homogenized by at least ten soil cores. Given that plots were structurally analogous, all samples followed by unified strict selection criteria including a slope of < 15°, the same soil type and a similar landform.

### Sampling, charcoal separation and physicochemical analysis

During sampling, the litter layer (a laminated mixing of small twigs, roots and fungal hyphae) was first removed and the organic layer (O horizon) comprising dark colored materials with fewer small roots and charcoal pieces were collected. Then, the top 15-cm mineral soil (A horizon) was collected using a soil auger (4 cm diameter). Soils were placed in the aseptic plastic bags for subsequent processing. Samples were transported to laboratory on ice, immediately sieved through a 2-mm mesh and stored at -80 °C (for DNA extraction) and 4 °C (for physicochemical analyses). Soil physicochemical properties included total C (%), N (%), P (mg kg^-1^), soil organic matter (SOM), pH, soil gravimetric water content, available P, dissolved organic C, dissolved inorganic C, dissolved organic N and exchangeable Na, K, Mg, and Ca. Detailed information on soil analyses is given in the Supplementary Information.

To minimize disturbance of microbial community, we adopted a direct hand-picked method (61, 62), rather than the water-floating method, to separate charcoal particles from the O-horizon soil. These “charcoal” particles in this study were defined as solid residual pieces derived from the pyrolysis of plant biomass by fire. Many researches showed that 1-3 mm char-particles (including biochar, charcoal) are easily extracted from soil (63–65), and in view of this phenomenon, we separated macroscopic charcoal particles (≥1 mm in diameter) from the sieved soil. We used a specific tweezer (N5, Dumont, Switzerland), which could isolate visible charcoal particles larger than 50 µm under microscope. Followed by the same criteria of microscopy and mineralogy, we collected charcoal approximately 15 g per plot for later analyses and characterization. The details of charcoal isolation from soil are presented in Supplementary Information. Collected charcoal pieces were crushed gently, homogenized and stored at -80 °C for DNA extraction and physicochemical analyses.

### Illumina sequencing and bioinformatic analysis

DNA was extracted from all soil (0.50 g) and charcoal (0.2-0.4 g) samples using the MP FastDNA SPIN Kit for soil (MP Biomedicals, Solon, OH, USA) according to the manufacturer’s protocol. Extracted DNA was stored at -20°C for a maximum of one week until amplicon library preparation began. PCR amplification was performed on the V4-V5 region of bacterial 16S rRNA gene using primers 515F/907R (66, 67) and the V4 region of 18S rRNA using primers 528F/706R (68, 69). The concentration and purity of DNA were monitored on 1% agarose gels. According to actual concentration, DNA was diluted to about 1 ng μL^-1^ using sterile water. 16S /18S rRNA genes were amplified used specific primers with the barcode. All PCR reactions were carried out in a volume of 30 μL using 15 μL of Phusion^®^ High-Fidelity PCR Master Mix (New England Biolabs), 0.2 μM of forward primers, 0.2 μM of reverse primers and 10 μL sample DNA. The PCR consisted of 98 °C for 1 min, then 30 cycles at 98°C for 10 s, 50 °C for 30 s, 72 °C for 30 s, and finally 72 °C for 5 min. PCR products were mixed in equidensity ratios and purified before sequencing libraries were generated. The library quality was assessed on the Qubit@ 2.0 Fluorometer (Thermo Scientific) and Agilent Bioanalyzer 2100 system. At last, the library was sequenced on an Illumina HiSeq 2500 and 250 bp paired-end reads were generated.

Briefly, the raw sequences from HiSeq platform were merged by using FLASH (70) and were assigned to each sample according to the specific barcodes. The sequences were pre-filtered and removed chimeras (71) by QIIME’s (72) quality filters. All effective sequences were then clustered into operational taxonomic units (OTUs) by Uparse (73) based on 97% similarity level. We picked the highest frequent sequence for each OTU as their representative sequences. We used the Ribosomal Database Project’s classifier (74) against the SILVA 132 database (75) to annotate taxonomic information for bacteria (16S) and eukaryotes (18S). RDP classifier bootstrap confidence values were 0.8-1. We normalized the read counts in OTU table by rarefying to the minimum sequence number within all samples. Moreover, to parse fungal OTUs by ecological guilds, we used FUNGuild to annotate fungal taxa into different functional groups (i.e. saprotroph, pathotroph and symbiotroph) (76). Raw sequence data were deposited into the Genome Sequence Archive (GSA) database under accession numbers CRA002389 that are publicly accessible.

### Statistical analysis

All statistical analyses were performed using R v. 3.5.3 (77) using the rarefied data on the following packages: phyloseq (78), ggplot2 (79), reshape2 (80), plyr (81), except for the analysis using non-rarefied data (82). Here, *α*-diversity metrics included Shannon index and OTU richness (namely species richness), which were calculated using the “plot_richness” function (phyloseq) and Picante package (83). The vegan package (84) was used to assess β-diversity via non-metric multidimensional scaling (NMDS) ordination using “metaMDS” function, based on Bray-Curtis dissimilarities. Bacterial and fungal communities in charcoal and soil samples were determined separately by the differential abundance analysis compared with the unburnt soils in each soil horizon. The package ‘DESeq2’ (85) was used to calculate the differential abundance (Log_2_ fold change in relative abundance of OTUs) using non-rarefied data. DESeq2 was run using the Wald test and an alpha of 0.01.

To reduce network complexity, we removed OTUs with relative abundances less than 0.001% of the total number of bacterial and fungal sequences, respectively. We then calculated all possible Spearman’s rank correlations between OTUs more than three sequences using the WGCNA package (86). The nodes in the networks represent microbial taxa (OTUs) and the edges represent significant correlations between the nodes. The *p* values were adjusted using the the Benjamini-Hochberg false discovery rate (FDR) controlling procedure (87), as implemented in the multtest package (88). A valid co-occurrence network was considered a statistically robust correlation between taxa when the correlation coefficients threshold was above 0.8 and FDR-adjusted *p* value was below 0.01. Network visualization was conducted using Cytoscape version 3.8.0 (89), and topographical properties (including average degree, average path length, network diameter, and clustering coefficient) of networks were calculated using Gephi software (90).

To investigate the importance of fire effect and soil properties, a permutational multivariate analysis of variance (PERMANOVA) was performed on Bray-Curtis dissimilarities using the “adonis” function of vegan (n_perm_=9999). Partial Mantel tests were conducted to test the effects of soil properties on the compositional similarity of OTUs using vegan package. Moreover, the effects of spatial variation (i.e. longitude and latitude) were controlled while testing the correlation of soil factors and the microbial community of burnt and unburnt soils. Soil properties, the relative abundances of taxa and *α*-diversity indices were performed by ANOVA, and pair-wise differences were assessed by Tukey’s HSD post-hoc tests as implemented in agricolae package. The Kruskal-Wallis test instead was used for the data with abnormal distribution or homoscedastic. To further test the effect of fire (including TSF and charcoal) on soil factors and soil community structure of bacteria and fungi, we built Structural Equation Modeling (SEM) by “sem” function from the lavaan package. A priori models based on the results of linear mixed-effect model (Fig. S7). We used fire effect (TSF) and fundamental edaphic factors (pH, dissolved organic C) as exogenous variables, and soil total N and P, and microbial community in charcoal as endogenous variables (Fig. 6). Exploratory analyses showed dissolved organic C had a better fit and explanation than soil organic matter or total C. Importantly, the microbial community colonized on the separated charcoal particles were hypothesized as key links between fire effect and soil microbial community. The goodness of fit was considered on the basis of chi-square test and root-mean-square error of approximation (RMSEA). A model could be accepted when *p*-value > 0.05, comparative fit index (CFI) > 0.95 and RMSEA < 0.05.

## Acknowledgements

This work was financially supported by the National Natural Science Foundation of China (41520104001, 41721001) and the 111 Project (B17039).

## Supplementary Information

### Supplementary Methods

*Additional site information*

*Soil analyses*

*Charcoal hand-picked method*

*References for Supplementary Methods*

### Supplementary Tables

**Table S1**. Field site information of fire-affected and unaffected soils.

**Table S2**. Effects of fire history on soil properties in soil O and A horizons.

**Table S3**. Effects of fire history on the relative abundances (specific OTUs × 100 / total number) of bacterial and fungal phyla in soil O and A horizons.

**Table S4**. Results from permutational multivariate analysis of variance with Bray-Curtis dissimilarity from 82 soil samples.

**Table S5**. Differential bacterial OTUs in the recent fires (RF) compared with unfired sites in O and A horizons.

**Table S6**. Differential fungal OTUs in the recent fires (RF), middle-term fires (MF), old fires (OF) compared with unfired sites in O and A horizons.

**Table S7**. Topological properties of networks obtained at the different fire histories and different horizons (O and A).

**Table S8**. Bacterial and fungal community structure association with soil factors by Partial Mental test in different fire histories and different horizons (O and A).

### Supplementary Figures

**Figure S1**. The study region of 44 sites where samples were collected in the Chinese boreal forest.

**Figure S2**. Soil properties for unburnt and burnt groups.

**Figure S3**. Non-metric multidimensional scaling ordination plot based on Bray-Curtis dissimilarity showing changes in soil bacterial (a) and fungal (b) community structures.

**Figure S4**. All samples using NMDS ordination based on Bray-Curtis dissimilarity to depict fire legacy effect on microbial communities.

**Figure S5**. The bacterial co-occurrence networks under recent fires (RF, b), middle-term fires (MF, c), old fires (OF, d) and fire-unaffected groups (UF, a) in A horizon.

**Figure S6**. The fungal co-occurrence networks under recent fires (RF, b), middle-term fires (MF, c), old fires (OF, d) and fire-unaffected groups (UF, a) in A horizon.

**Figure S7**. The general structure of *a priori* structural equation model including all possible pathways (1-17).

## References

1. Bowman DMJS, Balch JK, Artaxo P, Bond WJ, Carlson JM, Cochrane MA, D’Antonio CM, DeFries RS, Doyle JC, Harrison SP, Johnston FH, Keeley JE, Krawchuk MA, Kull CA, Marston JB, Moritz MA, Prentice IC, Roos CI, Scott AC, Swetnam TW, van der Werf GR, Pyne SJ. 2009. Fire in the Earth System. Science 324:481–484.

2. Bond W, Keeley J. 2005. Fire as a global ‘herbivore’: the ecology and evolution of flammable ecosystems. Trends in Ecology & Evolution 20:387–394.

3. Certini G. 2005. Effects of fire on properties of forest soils: a review. Oecologia 143:1–10.

4. Dooley SR, Treseder KK. 2012. The effect of fire on microbial biomass: a meta-analysis of field studies. Biogeochemistry 109:49–61.

5. Luo Y, Yu Z, Zhang K, Xu J, Brookes PC. 2016. The properties and functions of biochars in forest ecosystems. J Soils Sediments 16:2005–2020.

6. Smith NR, Kishchuk BE, Mohn WW. 2008. Effects of Wildfire and Harvest Disturbances on Forest Soil Bacterial Communities. Applied and Environmental Microbiology 74:216–224.

7. Sun H, Santalahti M, Pumpanen J, Köster K, Berninger F, Raffaello T, Asiegbu FO, Heinonsalo J. 2016. Bacterial community structure and function shift across a northern boreal forest fire chronosequence. Scientific Reports 6.

8. Daniel RM, Cowan DA. 2000. Biomolecular stability and life at high temperatures. Cellular and Molecular Life Sciences 57:250–264.

9. Holden SR, Treseder KK. 2013. A meta-analysis of soil microbial biomass responses to forest disturbances. Frontiers in Microbiology 4.

10. Pérez-Valera E, Goberna M, Verdú M. 2019. Fire modulates ecosystem functioning through the phylogenetic structure of soil bacterial communities. Soil Biology and Biochemistry 129:80–89.

11. Taş N, Prestat E, McFarland JW, Wickland KP, Knight R, Berhe AA, Jorgenson T, Waldrop MP, Jansson JK. 2014. Impact of fire on active layer and permafrost microbial communities and metagenomes in an upland Alaskan boreal forest. The ISME Journal 8:1904–1919.

12. Randerson JT, Liu H, Flanner MG, Chambers SD, Jin Y, Hess PG, Pfister G, Mack MC, Treseder KK, Welp LR, Chapin FS, Harden JW, Goulden ML, Lyons E, Neff JC, Schuur EAG, Zender CS. 2006. The Impact of Boreal Forest Fire on Climate Warming. Science 314:1130–1132.

13. Abatzoglou JT, Williams AP, Boschetti L, Zubkova M, Kolden CA. 2018. Global patterns of interannual climate-fire relationships. Global Change Biology 24:5164–5175.

14. Liu Z, Yang J, Chang Y, Weisberg PJ, He HS. 2012. Spatial patterns and drivers of fire occurrence and its future trend under climate change in a boreal forest of Northeast China. Global Change Biology 18:2041–2056.

15. van Mantgem PJ, Nesmith JCB, Keifer M, Knapp EE, Flint A, Flint L. 2013. Climatic stress increases forest fire severity across the western United States. Ecology Letters 16:1151–1156.

16. Shungart HH, Leemans Rik, Bonan GB. 1992. A Systems Analysis of the Global Boreal Forest. Cambridge University Press, Cambridge.

17. Xu H. 1998. Forests in Daxinganling Moutains China. Science Press, Beijing, China.

18. Hu H, Wei S, Sun L. 2012. Estimation of carbon emissions due to forest fire in Daxing’an Mountains from 1965 to 2010. Chinese Journal of Plant Ecology 36:629–644.

19. Chang Y, He HS, Hu Y, Bu R, Li X. 2008. Historic and current fire regimes in the Great Xing’an Mountains, northeastern China: Implications for long-term forest management. Forest Ecology and Management 254:445–453.

20. Wang X, He HS, Li X. 2007. The long-term effects of fire suppression and reforestation on a forest landscape in Northeastern China after a catastrophic wildfire. Landscape and Urban Planning 79:84–95.

21. Flannigan MD, Krawchuk MA, de Groot WJ, Wotton BM, Gowman LM. 2009. Implications of changing climate for global wildland fire. International Journal of Wildland Fire 18:483.

22. Santín C, Doerr SH, Kane ES, Masiello CA, Ohlson M, de la Rosa JM, Preston CM, Dittmar T. 2016. Towards a global assessment of pyrogenic carbon from vegetation fires. Global Change Biology 22:76–91.

23. Bird MI, Wynn JG, Saiz G, Wurster CM, McBeath A. 2015. The Pyrogenic Carbon Cycle. Annual Review of Earth and Planetary Sciences 43:273–298.

24. Preston CM, Schmidt MWI. 2006. Black (pyrogenic) carbon: a synthesis of current knowledge and uncertainties with special consideration of boreal regions. Biogeosciences 3:397–420.

25. Gavin DG, Hallett DJ, Hu FS, Lertzman KP, Prichard SJ, Brown KJ, Lynch JA, Bartlein P, Peterson DL. 2007. Forest fire and climate change in western North America: insights from sediment charcoal records. Frontiers in Ecology and the Environment 5:499–506.

26. Ohlson M, Dahlberg B, Økland T, Brown KJ, Halvorsen R. 2009. The charcoal carbon pool in boreal forest soils. Nature Geoscience 2:692–695.

27. Hart S, Luckai N. 2013. Charcoal function and management in boreal ecosystems. Journal of Applied Ecology 50:1197–1206.

28. Pietikainen J, Kiikkila O, Fritze H. 2000. Charcoal as a habitat for microbes and its effect on the microbial community of the underlying humus. Oikos 89:231–242.

29. Wardle DA, Zackrisson O, Nilsson M-C. 1998. The charcoal effect in Boreal forests: mechanisms and ecological consequences. Oecologia 115:419–426.

30. Bryanin S, Abramova E, Makoto K. 2018. Fire-derived charcoal might promote fine root decomposition in boreal forests. Soil Biology and Biochemistry 116:1–3.

31. Hestrin R, Torres-Rojas D, Dynes JJ, Hook JM, Regier TZ, Gillespie AW, Smernik RJ, Lehmann J. 2019. Fire-derived organic matter retains ammonia through covalent bond formation. Nature Communications 10.

32. Wardle DA, Nilsson M-C, Zackrisson O. 2008. Fire-derived charcoal causes loss of forest humus. Science 320:629–629.

33. Preston CM. 2009. Fire’s black legacy. Nature Geoscience 2:674–675.

34. Zimmermann M, Bird MI, Wurster C, Saiz G, Goodrick I, Barta J, Capek P, Santruckova H, Smernik R. 2012. Rapid degradation of pyrogenic carbon. Global Change Biology 18:3306–3316.

35. Butler OM, Elser JJ, Lewis T, Mackey B, Chen C. 2018. The phosphorus-rich signature of fire in the soil-plant system: a global meta-analysis. Ecology Letters 21:335–344.

36. Bowd EJ, Banks SC, Strong CL, Lindenmayer DB. 2019. Long-term impacts of wildfire and logging on forest soils. Nature Geoscience 12:113–118.

37. Cui X, Gao F, Song J, Sang Y, Sun J, Di X. 2014. Changes in soil total organic carbon after an experimental fire in a cold temperate coniferous forest: A sequenced monitoring approach. Geoderma 226–227:260–269.

38. Hu T, Sun L, Hu H, Weise DR, Guo F. 2017. Soil respiration of the Dahurian Larch (Larix gmelinii) forest and the response to fire disturbance in Da Xing’an Mountains, China. Scientific Reports 7:2967.

39. Blois JL, Williams JW, Fitzpatrick MC, Jackson ST, Ferrier S. 2013. Space can substitute for time in predicting climate-change effects on biodiversity. Proceedings of the National Academy of Sciences 110:9374–9379.

40. Carter Z, Sullivan B, Qualls R, Blank R, Schmidt C, Verburg P. 2018. Charcoal Increases Microbial Activity in Eastern Sierra Nevada Forest Soils. Forests 9:93.

41. Sun H, Santalahti M, Pumpanen J, Köster K, Berninger F, Raffaello T, Jumpponen A, Asiegbu FO, Heinonsalo J. 2015. Fungal community shifts in structure and function across a boreal forest fire chronosequence. Applied and Environmental Microbiology 81:7869–7880.

42. Cairney JWG, Bastias BA. 2007. Influences of fire on forest soil fungal communities. Can J For Res 37:207–215.

43. Holden SR, Gutierrez A, Treseder KK. 2013. Changes in Soil Fungal Communities, Extracellular Enzyme Activities, and Litter Decomposition Across a Fire Chronosequence in Alaskan Boreal Forests. Ecosystems 16:34–46.

44. Ferrenberg S, O’Neill SP, Knelman JE, Todd B, Duggan S, Bradley D, Robinson T, Schmidt SK, Townsend AR, Williams MW, Cleveland CC, Melbourne BA, Jiang L, Nemergut DR. 2013. Changes in assembly processes in soil bacterial communities following a wildfire disturbance. The ISME Journal 7:1102–1111.

45. Xiang X, Shi Y, Yang J, Kong J, Lin X, Zhang H, Zeng J, Chu H. 2015. Rapid recovery of soil bacterial communities after wildfire in a Chinese boreal forest. Scientific Reports 4.

46. Pérez-Valera E, Goberna M, Faust K, Raes J, García C, Verdú M. 2017. Fire modifies the phylogenetic structure of soil bacterial co-occurrence networks. Environmental Microbiology 19:317–327.

47. Paredes-Sabja D, Setlow P, Mahfuzur S. 2011. Germination of spores of Bacillales and Clostridiales species: mechanisms and proteins involved. Trends in Microbiology 19:85–94.

48. Sharples GP, Williams ST, Bradshaw RM. 1974. Spore formation in the Actinoplanaceae (Actinomycetales). Arch Microbiol 101:9–20.

49. Mikita-Barbato RA, Kelly JJ, Tate RL. 2015. Wildfire effects on the properties and microbial community structure of organic horizon soils in the New Jersey Pinelands. Soil Biology and Biochemistry 86:67–76.

50. Duhamel M, Wan J, Bogar LM, Segnitz RM, Duncritts NC, Peay KG. 2019. Plant selection initiates alternative successional trajectories in the soil microbial community after disturbance. Ecol Monogr 89.

51. Knelman JE, Graham EB, Trahan NA, Schmidt SK, Nemergut DR. 2015. Fire severity shapes plant colonization effects on bacterial community structure, microbial biomass, and soil enzyme activity in secondary succession of a burned forest. Soil Biology and Biochemistry 90:161–168.

52. Wang C, Wang G, Wang Y, Rafique R, Ma L, Hu L, Luo Y. 2016. Fire alters vegetation and soil microbial community in alpine meadow. Land Degradation & Development 27:1379–1390.

53. Dahlberg A. 2002. Effects of fire on ectomycorrhizal fungi in Fennoscandian boreal forests. Silva Fenn 36.

54. Glassman SI, Levine CR, DiRocco AM, Battles JJ, Bruns TD. 2016. Ectomycorrhizal fungal spore bank recovery after a severe forest fire: some like it hot. ISME J 10:1228–1239.

55. Oguntunde PG, Fosu M, Ajayi AE, van de Giesen N. 2004. Effects of charcoal production on maize yield, chemical properties and texture of soil. Biol Fertil Soils 39:295–299.

56. Yu M, Meng J, Yu L, Su W, Afzal M, Li Y, Brookes PC, Redmile-Gordon M, Luo Y, Xu J. 2019. Changes in nitrogen related functional genes along soil pH, C and nutrient gradients in the charosphere. Science of The Total Environment 650:626–632.

57. DeLuca TH, MacKenzie MD, Gundale MJ, Holben WE. 2006. Wildfire-Produced Charcoal Directly Influences Nitrogen Cycling in Ponderosa Pine Forests. Soil Science Society of America Journal 70:448.

58. Wang C, Gower ST, Wang Y, Zhao H, Yan P, Bond-Lamberty BP. 2001. The influence of fire on carbon distribution and net primary production of boreal Larix gmelinii forests in north-eastern China. Global Change Biology 7:719–730.

59. Oliveras I, Román-Cuesta RM, Urquiaga-Flores E, Quintano Loayza JA, Kala J, Huamán V, Lizárraga N, Sans G, Quispe K, Lopez E, Lopez D, Cuba Torres I, Enquist BJ, Malhi Y. 2018. Fire effects and ecological recovery pathways of tropical montane cloud forests along a time chronosequence. Global Change Biology 24:758–772.

60. Ludwig SM, Alexander HD, Kielland K, Mann PJ, Natali SM, Ruess RW. 2018. Fire severity effects on soil carbon and nutrients and microbial processes in a Siberian larch forest. Global Change Biology 24:5841–5852.

61. Carvalho LC d. S, Fearnside PM, Nascimento MT, Barbosa RI. 2018. Amazon soil charcoal: Pyrogenic carbon stock depends of ignition source distance and forest type in Roraima, Brazil. Global Change Biology 24:4122–4130.

62. Nguyen BT, Lehmann J, Kinyangi J, Smernik R, Riha SJ, Engelhard MH. 2008. Long-term black carbon dynamics in cultivated soil. Biogeochemistry 89:295–308.

63. Dai Z, Webster TM, Enders A, Hanley KL, Xu J, Thies JE, Lehmann J. 2017. DNA extraction efficiency from soil as affected by pyrolysis temperature and extractable organic carbon of high-ash biochar. Soil Biology and Biochemistry 115:129–136.

64. Lin Y, Munroe P, Joseph S, Kimber S, Van Zwieten L. 2012. Nanoscale organo-mineral reactions of biochars in ferrosol: an investigation using microscopy. Plant and Soil 357:369–380.

65. Ye J, Joseph SD, Ji M, Nielsen S, Mitchell DRG, Donne S, Horvat J, Wang J, Munroe P, Thomas T. 2017. Chemolithotrophic processes in the bacterial communities on the surface of mineral-enriched biochars. The ISME Journal 11:1087–1101.

66. Biddle JF, Fitz-Gibbon S, Schuster SC, Brenchley JE, House CH. 2008. Metagenomic signatures of the Peru Margin subseafloor biosphere show a genetically distinct environment. Proceedings of the National Academy of Sciences 105:10583–10588.

67. Stubner S. 2002. Enumeration of 16S rDNA of Desulfotomaculum lineage 1 in rice field soil by real-time PCR with SybrGreenk detection. Journal of Microbiological Methods 50:155–164.

68. Cheung MK, Au CH, Chu KH, Kwan HS, Wong CK. 2010. Composition and genetic diversity of picoeukaryotes in subtropical coastal waters as revealed by 454 pyrosequencing. The ISME Journal 4:1053–1059.

69. Elwood HJ. 1985. The small-subunit ribosomal RNA gene sequences from the Hypotrichous Ciliates Oxytriclha nova and Stylonychia pustulata. Molecular Biology and Evolution 2:399–410.

70. Magoc T, Salzberg SL. 2011. FLASH: fast length adjustment of short reads to improve genome assemblies. Bioinformatics 27:2957–2963.

71. Haas BJ, Gevers D, Earl AM, Feldgarden M, Ward DV, Giannoukos G, Ciulla D, Tabbaa D, Highlander SK, Sodergren E, Methe B, DeSantis TZ, The Human Microbiome Consortium, Petrosino JF, Knight R, Birren BW. 2011. Chimeric 16S rRNA sequence formation and detection in Sanger and 454-pyrosequenced PCR amplicons. Genome Research 21:494–504.

72. Caporaso JG, Kuczynski J, Stombaugh J, Bittinger K, Bushman FD, Costello EK, Fierer N, Peña AG, Goodrich JK, Gordon JI, Huttley GA, Kelley ST, Knights D, Koenig JE, Ley RE, Lozupone CA, McDonald D, Muegge BD, Pirrung M, Reeder J, Sevinsky JR, Turnbaugh PJ, Walters WA, Widmann J, Yatsunenko T, Zaneveld J, Knight R. 2010. QIIME allows analysis of high-throughput community sequencing data. Nature Methods 7:335–336.

73. Edgar RC. 2013. UPARSE: highly accurate OTU sequences from microbial amplicon reads. Nature Methods 10:996–998.

74. Wang Q, Garrity GM, Tiedje JM, Cole JR. 2007. Naive bayesian classifier for rapid assignment of rRNA sequences into the new bacterial taxonomy. Applied and Environmental Microbiology 73:5261–5267.

75. Quast C, Pruesse E, Yilmaz P, Gerken J, Schweer T, Yarza P, Peplies J, Glöckner FO. 2012. The SILVA ribosomal RNA gene database project: improved data processing and web-based tools. Nucleic Acids Research 41:D590–D596.

76. Nguyen NH, Song Z, Bates ST, Branco S, Tedersoo L, Menke J, Schilling JS, Kennedy PG. 2016. FUNGuild: An open annotation tool for parsing fungal community datasets by ecological guild. Fungal Ecology 20:241–248.

77. R Development Core Team. 2011. R: A language and environment for statistical computing. R foundation for statistical computing Vienna, Austria.

78. McMurdie PJ, Holmes S. 2013. phyloseq: An R Package for reproducible interactive analysis and graphics of microbiome census data. Plos one 8:e61217.

79. Wickham H. 2016. ggplot2. Springer International Publishing, Cham.

80. Wickham H. 2007. Reshaping data with the reshape Package. Journal of Statistical Software 21.

81. Wickham H. 2011. The split-apply-combine strategy for data analysis. Journal of Statistical Software 40.

82. McMurdie PJ, Holmes S. 2014. Waste not, want not: why rarefying microbiome data Is inadmissible. PLoS Comput Biol 10:e1003531.

83. Kembel SW, Cowan PD, Helmus MR, Cornwell WK, Morlon H, Ackerly DD, Blomberg SP, Webb CO. 2010. Picante: R tools for integrating phylogenies and ecology. Bioinformatics 26:1463–1464.

84. Oksanen J. 2007. vegan?: Community Ecology Package. R package version 1.8-5.

85. Love MI, Huber W, Anders S. 2014. Moderated estimation of fold change and dispersion for RNA-seq data with DESeq2. Genome Biol 15:550.

86. Langfelder P, Horvath S. 2008. WGCNA: an R package for weighted correlation network analysis. BMC Bioinformatics 9:559.

87. Benjamini Y, Krieger AM, Yekutieli D. 2006. Adaptive linear step-up procedures that control the false discovery rate. Biometrika 93:491–507.

88. Pollard KS, Dudoit S, van der Laan MJ. 2005. Multiple Testing Procedures: the multtest Package and Applications to Genomics, p. 249–271. In Gentleman, R, Carey, VJ, Huber, W, Irizarry, RA, Dudoit, S (eds.), Bioinformatics and Computational Biology Solutions Using R and Bioconductor. Springer New York, New York, NY.

89. Shannon P, Markiel A, Ozier O, Baliga NS, Wang JT, Ramage D, Nada A, Schwikowski B, ldeker T. 2003. Cytoscape: a software environment for integrated models of biomolecular interaction network. Genome Research 13:2498–2504.

90. Bastian M, Heymann S, Jacomy M. 2009. Gephi?: an open source software for exploring and manipulating networks.

